# A phylogenetic host range index reveals contrasted relationships between phage virulence and specialisation

**DOI:** 10.1101/2024.05.07.592886

**Authors:** Clara Torres-Barceló, Claudine Boyer, Julian R. Garneau, Stéphane Poussier, Isabelle Robène, Benoit Moury

**Affiliations:** INRAE, Pathologie Végétale, F-84140, Montfavet, France; CIRAD, UMR PVBMT, F-97410 St Pierre, La Réunion, France; Department of Fundamental Microbiology, University of Lausanne, CH-1015 Lausanne, Switzerland; Université de la Réunion, UMR PVBMT, F-97410 St Pierre, La Réunion, France

**Keywords:** phage, host range, virulence, CRISPR, *Ralstonia solanacearum*, plant pathogen, epidemiology

## Abstract

Phages are typically known for having a limited host range, targeting various strains within a specific bacterial species. However, factors like the phylogeny or epidemiology of host bacteria are often disregarded, despite their potential influence on phage specialization and virulence. This research utilizes a new “phylogenetic host range index” that accounts for the genetic diversity of bacterial hosts, to classify phages into specialists and generalists accurately. We provide evidence that the CRISPR-Cas immune system of bacteria more frequently targets generalist phages than specialist phages. We explore the hypothesis that generalist phages might exhibit lower virulence than specialist ones due to potential evolutionary trade-offs between host range breadth and virulence. Importantly, contrasted correlations between phage virulence and host range depend on the epidemiological context. A trade-off was confirmed in a homogeneous bacterial epidemiology situation, but not in more complex epidemiological scenario, where no apparent costs were detected for phages adapted to a wide range of hosts. This study highlights the need for genetic analyses in phage host range and of investigating ecological trade-offs that could improve their applications in biocontrol or therapy.

## INTRODUCTION

Current studies propose that the great majority of known bacteriophages (phages), the viruses of bacteria, possess a narrow host range, typically being able to replicate in some strains of one species, and rarely in several closely related species (1). Compared to animal or plant pests and pathogens, genomic analyses are insufficiently implemented when studying the host range of phages and only include phenotypic data (2–4). This has consequences for the understanding of the association of phage host range with bacterial epidemiology or the evolutionary ecology of phage-bacteria interactions, including host jumps. For example, it is not clear if phage specialization is restricted by the geographical origin or by the phylogenetic patterns of their hosts. As specialization is a common trait in pathogens, it could theoretically impose a trade-off on fitness and virulence in different hosts, for instance, as a consequence of the difficulty to optimise viral infection on very diverse hosts (5). The negative correlation between niche expansion and virulence has been proven in parasitic wasps and aphids, microsporidians and shrimps, or herbivorous mites and plants, among others, but we know little in phages and bacteria (6–9).

To assess the host range of phages is a complex aim, with differing methods, analyses, and numerous bacterial strains to be potentially included as hosts in the experiments. In the past, most studies used only qualitative spot tests (recording the occurrence or absence of bacterial lysis by the phage in a bacterial lawn) that did not account for abortive infections (a resistance mechanism of bacteria attacked by phages that impedes the production of phage progeny) (10). While some recent works use quantitative tests, they often lack statistical analyses specifically adapted to assess phage host range. This hinders comparisons between studies, due to the different panels of bacteria and phages tested and the vast microbial genetic diversity. Defining the specialisation or generalism of phages remains a subjective matter that would benefit from more precise and standardized approaches.

A key analysis of phage’s host range is to assess the structure of the host-parasite interaction matrix (11–13). Depending on the genetic, evolutionary and mechanistic patterns of host-parasite interactions, contrasted scores for nestedness and modularity are expected. Nestedness measures the tendency of hosts and parasites to have a hierarchical organization in terms of resistance spectra and host ranges, respectively. In contrast, modularity measures the strength by which a matrix can be subdivided into a several of groups (i.e. modules) of hosts and parasites characterized by successful infections, while infections are rarer or less efficient between hosts and parasites that belong to different modules. Recently, statistically robust analyses of quantitative assessments of nestedness and modularity in network-based analyses have been optimized and allow a better interpretation of the phage-bacteria interactions (14). This approach is helpful to disentangle phage-bacteria interactions, but nestedness or modularity values refer to a set of phages and bacteria rather than to an individual phage. They also ignore the phylogenetic relatedness of the bacterial strains used, with some studies including strains of a bacterial species and others different potential host species or even genera (15). In fact, matrix analysis of phage-bacteria interactions have often resulted in nested patterns probably due to the use of non-genetically diverse bacteria or phages (11, 16, 17). Modularity has been recently observed when the evolutionary or ecological scales considered between bacteria and or phages are wide (18, 19). New methods to evaluate phage host range integrating the vast amount of sequencing data currently available are timely and feasible.

The model bacteria used in this study are strains from the *Ralstonia solanacearum* species complex (RSSC or *Ralstonia* spp.). They are responsible of bacterial wilt, one of the major bacterial diseases of plants. These pathogenic bacteria can infect all the solanaceous plant family, which includes tomato, potato and aubergine, and 23 more botanical families, such as the Musaceae (banana). In tropical areas it can be devastating, affecting essential crops and threatening food security of entire countries (20, 21). The RSSC has long been known for its high genetic and phenotypic diversity (22). Recently, RSSC strains were classified into three species and four major phylogenetic groups, named phylotypes: *R. pseudosolanacearum* including phylotypes I and III strains, *R. solanacearum* grouping phylotypes IIA and IIB strains, and *R. syzygii* comprising phylotypes IV strains (23, 24). Each phylotype is subdivided into many sequevars based on partial sequencing of the endoglucanase (*egl*) gene (25). Moreover, each sequevar comprises several haplotypes based on a multilocus variable-number of tandem repeats (26).

*Ralstonia spp.* are distributed worldwide, with the South-West Indian Ocean (SWIO) being one of the best studied areas. This work concerns the SWIO islands Reunion and Mauritius. The diversity of the RSSC strains in Reunion Island was first described in 1993 and phylotypes I, IIB-1 and III were reported (27). A recent survey revealed that 70% of the strains belonged to phylotype I, with sequevar I-31 being the most prevalent, accounting for 94% of all strains among the identified sequevars (I-13, I-31, I-33, IIA-36, IIB-1, and III-19) (28). In Mauritius, phylotype I was described first (29). Later, three potato brown rot epidemics (2005, 2006, and 2008) caused by sequevar IIB-1 were reported (30, 31). However, more recently, only phylotype I (with the exception of two phylotype IV strains) was disclosed, with a broad phylogenetic diversity including 5 sequevars (I- 14, I-15, I-18, I-31, and I-33), and a lower prevalence of sequevar I-31 (19% of phylotype I) than in Reunion (28, 32). In short, the present scenario regarding the genetic diversity of phylotype I, dominant in the SWIO, is contrasting between the two islands, with one sequevar almost exclusive in Reunion and a huge diversity of sequevars in Mauritius (32).

Despite its genetic diversity and worldwide distribution, the interactions of RSSC with phages remain largely uncovered. Only around 50 phage genomes attacking *Ralstonia* spp. have been published (NCBI last accessed August 2023), although many more phages have been isolated. Most studies focus on the characterisation of one or few phages and their biocontrol potential, showing promising results (33, 34). The host range of these phages, as well as their capacity to target all the genetic and geographical variability of RSSC, are unknown. Previously described genetically diverse RSSC phages isolated in Mauritius and Reunion Islands, together with the wide genetic diversity of their host, provide us with a unique opportunity to test ecological and evolutionary hypotheses on phage host range structure and evolution, as well as to implement a new host range analysis methodology (35).

An important element related to specialisation are the defence systems of bacteria and CRISPR-Cas in particular. These systems are molecular records of the environmental conditions linked to viral abundance and diversity (36). Even if CRISPR-Cas systems have not been proved to be the primary anti-phage defence system in *Ralstonia* spp., 31% of the genomes harbour one (37). Building on this, our hypothesis proposes a correlation between CRISPR-Cas spacer acquisitions and the host range of phages. Specifically, we anticipate that generalist phages will be more frequently targeted by this immune system compared to specialist phages.

## MATERIAL AND METHODS

### Phages

The 23 *Ralstonia* spp. phages used in this study have been described elsewhere (35). Phage genomes GenBank accession numbers are MT740725 to MT740747. The strain RUN3665 (sequevar I-31) was used as the common host for subsequent phage amplification and titer standardisation of Reunion Island phages. Mauritian phages were amplified and standardized in three different hosts strains, because no common host could be found in a host range pre-screening analysis: RUN4407 (sequevar I-15), RUN4833 (sequevar I-33) and RUN5163 (sequevar I-31). All phages were sequenced after the amplification.

### Bacteria

All inoculations and bacterial cultures were carried out in semi-selective Kelman culture medium containing triphenyltetrazolium chloride. Luria Bertani medium was used for non-RSSC bacteria. All bacterial cultures were incubated at 28°C and liquid ones were agitated at 80 or 120 rpm, when including phages or not, respectively.

### Host range assay

RSSC strains and three strains of closely related species (*Burkholderia cepacia*, *Ralstonia eutropha* and *Ralstonia pickettii),* belonging to the collection of the “Pôle de Protection des Plantes” in Reunion Island, were used for this study (Table S1). The strain phylotypes and sequevars were selected to be representative of the specific genetic diversity of RSSC in Reunion and Mauritius, in addition to represent diversity in the SWIO region and the world. At least five strains per phylotype were used. The ten Reunion phages were exposed to 52 RSSC and the 13 Mauritian phages to 63 RSSC, in addition to the three non-RSSC strains.

The titer of phages was homogenized to 3.5x10⁵ pfu/mL for Mauritius, and to 1x10⁶ pfu/mL for Reunion, due to technical reasons. Drops of 10 µL of several ten-fold serial dilutions were deposited in triplicate on the surface of the semi-solid bacterial lawn of each strain. Phage fitness or the number of phages able to replicate on each strain (pfu/mL) was recorded using the most appropriate dilution.

### Phage phylogenetic host range index

A phylogenetic host range index (PHRI) of each phage was calculated, as a proxy of the diversity of bacterial strains that it can attack and how well it replicates in them. It accounts for the number of bacterial strains targeted, the relative phage replication (pfu/mL) on each strain (i.e. evenness), and the phylogenetic distance between bacterial hosts (38, 39). The PHRI index is based on phylogenetic abundance-based Hill numbers that incorporates relative abundance and species richness, as calculated with the “hilldiv” R package. The data used for this analysis were the quantitative matrix of phage-bacteria host range interactions (phage fitness on each strain) and a phylogenetic tree of the bacterial strains tested. The phylogenetic distance between bacterial hosts was calculated by using the phylogenetic tree of the RSSC endoglucanase (*egl*) gene, as previously established to study RSSC diversity (25). Due to the different RSSC strains used to standardise phage titer of Mauritius and Reunion strains, the PHRI was calculated separately for phages of each geographical origin.

### Genomic analyses

All phages were re-annotated using PROKKA (40), with a priority annotation using the PHASTER protein database (version Dec 22, 2020) (41). To establish the list of shared and unique proteins between each pair of phages, the predicted phage proteomes were analysed pairwise using Roary at 90% protein identity threshold (42). The tables for the proteins’ presence-absence comparisons can be found in Table S2.

All phage genomes were submitted to CRISPRDetect (43) to search for potential genes, repeats or spacers. Default parameters were used for the first scan (CRISPR likelihood score >= 2.50), and permissive parameter CRISPR likelihood score >= 0.5 for the second scan. All phage genomes were also submitted to CRISPRCasTyper with suggested default parameters (44). Then, all 106 CRISPR spacers from available RSSC genomes were downloaded from the CRISPI database (45). The known CRISPR spacers from *R. solanacearum* were then submitted to CRISPERTarget (46).

### Phage-bacteria network structure analysis

The nestedness and modularity of the two phage-bacteria matrices (Mauritius and Reunion) were estimated, and their statistical significance tested respectively with the ‘bipartite’ and ‘igraph’ packages of R software as described in Moury *et al*. (14). Only the subset of bacterial strains where at least one phage could replicate was used in this analysis. Prior to these analyses, in each matrix separately, infection values (titer of each phage measured on each bacterial strain) were binned into ten intervals of equal size, as described in Moury *et al*. (14). More details can be found as Supplementary Methods.

### Phage host range phylogenetic signal

A host range phylogenetic signal of each phage was calculated, to test the dependence in targeting phylogenetically-related hosts. We used the same matrix data of phage fitness and the same bacterial phylogenetic trees as for the PHRI calculation and applied the “MCMCglmm” R package.

### Phage virulence

The virulence of ten Reunion phages and eleven Mauritian ones (Hennie and Hyacinthe phages were not included due to their extreme specificity) was measured by their inhibitory effect on bacterial growth. The three most abundant bacterial strains on each island were used for each set of phages, namely RUN4407 (sequevar I-15), RUN4833 (sequevar I-33) and RUN5163 (sequevar I- 31) for Mauritius phages, and different haplotypes of sequevar I-31 (RUN3665, RUN3014, RUN3692) for Reunion phages. First, exponential-phase bacteria were incubated overnight and the optical density at 600 nm (OD600) homogenized to 0.1 (10⁸ cfu/mL) in the morning. Phages were tested against bacteria at 10^6^ pfu/mL, corresponding to a phage:bacteria ratio (MOI or Multiplicity Of Infection) of 0.01, in a final volume of 200 μL. The chosen phage dose represented an intermediate dose capable of reducing bacterial growth substantially but not completely. Three replicates of growth curves were recorded by measuring the OD600 every 15 minutes over 24 hours. All experiments were done using the plate spectrophotometer Bioscreen (Oy Growth Curves Ab Ltd., Finland). Positive control (phage-free bacterial cultures) and negative ones (bacteria-free) were included in each plate.

To analyse the data, an average of all OD600 measurements over time of each replicate culture was calculated. The mean blank value (negative control without bacteria and phages) was subtracted from all the mean OD600 values. Replicates of the positive controls (N=36) showed no significant difference between plates (days) (GLM p = 0.09, t-value = 1.73), therefore all data were analysed together. The inhibitory effect of phages on bacterial growth was calculated as the percent average growth difference between treatments and positive controls:

Phage virulence =100-((phage assay*100)/positive control)

### Correlation analysis between phage fitness and host range

For a given phage, we estimated the average fitness as the average of the fitness values measured on the bacterial strains for which replication was higher than zero (i.e. non sensitive host bacterial strains were discounted from the calculation). As the calculation of the phage PHRI is partly based on their fitness estimates, there is a risk of spurious correlation when analysing the correlation between the non-independent ‘PHRI’ and ‘fitness’ variables (47). To circumvent this pitfall, we defined an *ad hoc* null hypothesis based on the distribution of the coefficients of correlation obtained by using 1,000 random permutations, as suggested previously (48). For each permutation, the phage fitness values were first randomly permutated across bacterial strains. Then, the PHRI was calculated based on this permutated dataset and the Pearson’s correlation coefficient between PHRI and fitness was calculated. The comparison between the distribution of the 1,000 Pearson’s correlation coefficients obtained from random permutations and the correlation coefficient from the actual data was used to test the linear dependence between the fitness and the PHRI among phages.

### Additional statistical analyses

A non-parametric Kruskal-Wallis test was performed to test the significance of the links between phages’ PHRI, host range phylogenetic signal, virulence or fitness as dependent variables, on the following explanatory variables: taxonomy (species and genus), life style (temperate *versus* virulent), and geographical origin (island).

A Spearman correlation analysis was used to test the association of the phages’ PHRI, host range phylogenetic signal, virulence or fitness, and the following variables: genome size, GC % and protein content (35).

A Levene’s test for homogeneity of variances was done to compare the PHRI of the two islands. A non-parametric Kruskall-Wallis or ANOVA tests were used to analyse the effect of phage virulence on each bacterial strain. The Tukey’s *post-hoc* test of multiple comparisons was used to determine significant differences between the means of variables that have more than two categories. All statistical analyses were performed using R version 3.5.1 (http://cran.r-project.org/). Model assumptions were tested by visually exploring model residuals.

## RESULTS AND DISCUSSION

### 1. The host range of *Ralstonia* spp. phages extends from extreme specialists to wide generalists

Considering the high genetic diversity of the RSSC and its diverse epidemiology, we examined if phages were adapted only to the local diversity of RSSC or was extended beyond. To answer this question, we confronted thirteen Mauritius and ten Reunion phages to local and international strains of the host species. None of the phages targeted the three non-RSSC species tested, proving the restricted host range of phages beyond the species complex. All of them targeted strains from phylotype I, the most prevalent (87%) in the SWIO, reflecting their adaptation to the local epidemiology (28) (Fig. 1A and B). Eleven phages (7/10 from Reunion, 4/13 from Mauritius) replicated only on strains belonging to this phylotype, and 12 others (3/10 from Reunion, 9/13 from Mauritius) replicated in strains from this phylotype as well as from phylotypes II and IV (Fig. 1). Two RSSC strains from phylotype IV were isolated in 2017 in Mauritius, but phylotype IV has never been recorded in Reunion (28). The replication of phages from both islands on phylotype IV bacteria is probably explained by a genetic convergence of phage replication mechanisms or by the existence of conserved receptors in bacteria from phylotypes IV and II, and/or phylotypes IV and I. No phage could replicate in the tested strains from phylotype III, represented only in Reunion at a low frequency (8%) and on rare plant hosts (28). No strains from phylotype II have been recorded in Mauritius since 2010 but they are currently present in Reunion (22%). These results suggest that the epidemiological history of bacteria in the islands could be reflected, at least partially, in the phages host range, with many Mauritian phages preserving their capacity to target the now disappeared phylotype II.

**Fig. 1.**
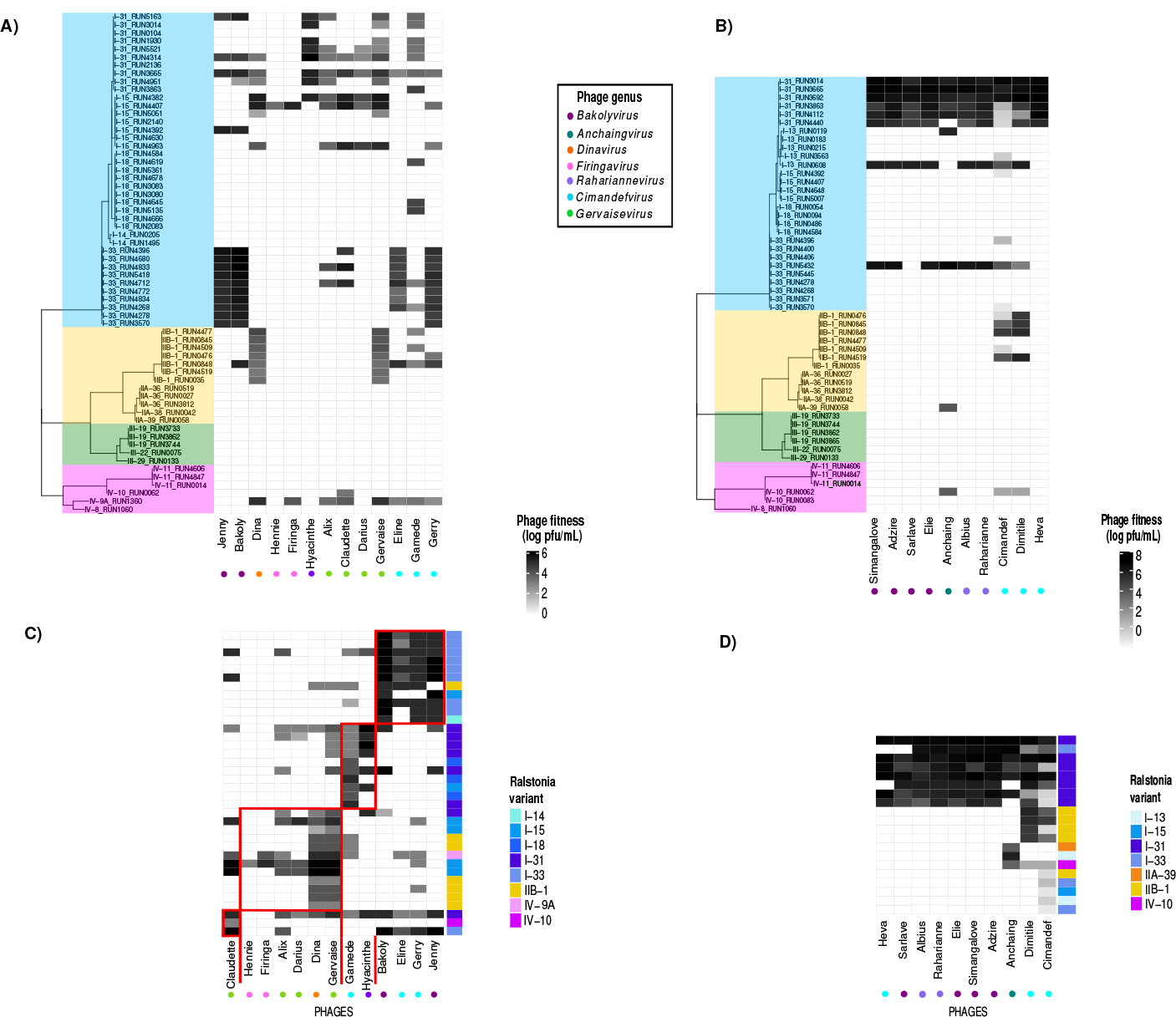
Quantitative heatmaps representing the host range of phages. The amount of progeny (pfu/mL) produced by each phage on each bacterial strain is indicated (grey gradient code), phage genera are detailed at the bottom with colour dots, and bacterial phylotypes are indicated in different colours. Panels A and B show phylogenetic heatmaps of 63 RSSC strains against 13 Mauritian phages, and 52 RSSC strains against ten Reunion phages, respectively. Panels C and D illustrate quantitative heatmaps with the network analysis of the host range of phages (only positive phage-bacteria interactions). The modular structure of Mauritius, with the four modules (C), and the nested structure in Reunion (D) phage-bacteria interactions, can be distinguished.

At the sequevar level, all phages but two Mauritian ones (Hennie and Firinga) target at least one strain of sequevar I-31, the most dominant genetic variant of the RSSC in Reunion (Fig. 1A). Seven out of ten Reunion phages target only strains of sequevar I-31, suggesting a strong specialisation (Fig. 1B). Most Mauritian phages target a relatively wide diversity of phylotype I sequevars, namely I-14, I-15 and I-33, besides I-31. The overall replication (pfu/mL) of phages of Reunion in the tested strains was higher than that of Mauritius phages (χ2 = 12.062, df = 1, p ˂ 0.001), suggesting that the fitness of Mauritius phages could be reduced as a trade-off of their generalist host range, even considering initial titer variations between Mauritian and Reunion phage preparations.

We then included in the analysis the genetic distance between the tested bacteria, in order to obtain a quantitative and more precise measure of phage host range. This is especially relevant in our dataset to avoid the bias of overrepresented phylotype I strains. We used a phylogenetic diversity index based on Allen’ H that accounts for the number of bacterial strains targeted (i.e. infected), the relative phage replication (pfu/mL) on each strain (i.e. evenness), and the phylogenetic distance between bacterial hosts. The phylogenetic host range index (PHRI) was significantly higher for Mauritian than Reunion phages (χ2 = 11.635, df = 1, p ˂ 0.001), meaning that they can target and replicate better on more strains that are genetically more distant (Fig. 2A). The PHRI of Mauritian phages has also a larger variation than that of Reunion phages (Levene’s Test F_1,21_ = 4.3355, p = 0.0497), ranging from extreme specialist (Hennie) to generalist phages (Bakoly and Gamede) (Fig. 2A). The PHRI of Reunion phages is far more homogeneous and narrower, reflecting the specialisation of most of them on closely related strains from sequevar I-31.

**Fig. 2.**
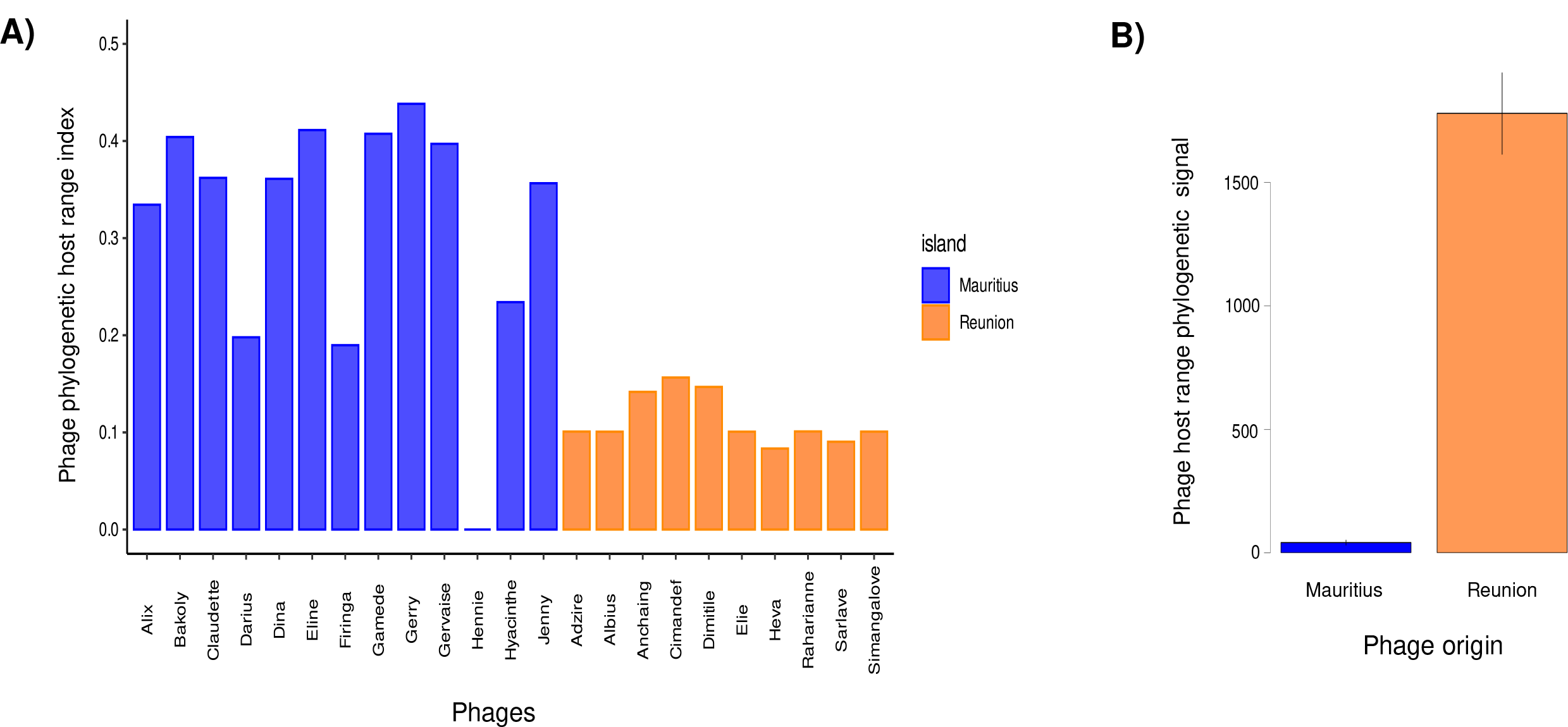
(A) Phage pylogenetic diversity index related to Hill numbers and (B) Phylogenetic signal of the strains targeted by *Ralstonia* spp. phages, relative to the island of origin of each phage. Data are average phylogenetic signal and standard errors. Both datasets represent 13 Mauritian phages and 33 RSSC strains and ten Reunion phages, and 20 RSSC strains.

The larger host range breadth and diversity of Mauritian phages is likely associated with the more complex epidemiological diversity of RSSC in the island, with a previously calculated Simpson’s Diversity Index of 0.907 and a genetic richness of 16.35, compared to the 0.485 and 5.41 values of bacteria isolated from Reunion (32). This can be partially explained by different agricultural practices. In Mauritius, there are local potato and tomato breeding practices that may mix and spread the RSSC strains. Until now, certified seeds have been imported in Reunion from Metropolitan France. It is important to highlight a potential bias in phage sampling for Reunion. Our focus was on the agriculturally rich, low-altitude regions of the island, where phylotype I is more prevalent and to which phages are shown to be adapted. At high altitude areas of Reunion, more phages adapted to phylotype II, and even III, would probably had been found, because of the lower temperature optimum and higher frequency of this RSSC variant in potato crops (49). This spatial compartmentalization does not exist in Mauritius. In summary, we quantified the host range of phages integrating the host genetic information, and observed important differences related to their geographical origin and bacterial epidemiology. Phages host range is adapted to the local epidemiology of bacteria but some generalist phages are able to target distantly-related strains.

### 2. Bacterial genomic analyses confirm the widespread occurrence of generalist phages

The genomic novelty of these phages limits the understanding of the encoded gene functions and their association with the host range analysis. In fact, the annotation of the genomes of all 23 viruses is partial, with more than 60% of proteins having unknown functions, and only few tail fibre genes clearly determined (which usually contain the receptor-binding proteins). We compared the genome of six pairs of phage species from the same genus (Table S2) with very different host ranges (i.e. Heva vs Cimandef, two species from the genus Cimandefvirus that are the least and most generalist phages of Reunion, respectively) using several up-to-date methods. The analysis did not reveal radical differences in gene content or gene length, except the presence of more annotated endonucleases in phages with narrower host range (Table S2). For instance, phages Heva and Simangalove, with a narrow host range, possess a CRISPR-associated endonuclease Cas9. Further bioinformatic and experimental analyses would be necessary to understand these findings.

Regarding bacterial genomes, bioinformatic analysis revealed that from the 116 CRISPR-Cas spacers published within RSSC genomes, 36 could target many generalist phages in our collection. Specifically, 19 hits were detected for Bakoly, 13 for Eline, and one for Darius, Gamede, Gervaise and Heva. All but Heva are phages isolated in Mauritius and categorized as generalist phages according to the PHRI values. These results suggest that these phages, particularly Bakoly and Eline, have been repeatedly targeted by the CRISPR-Cas system. Therefore, our data support the hypothesis suggesting this immune system acts more frequently against generalist phages compared to specialist phages, as the former encounter their hosts more often.

Unfortunately, the RSSC genome sequences used in this study are not available. A previous study showed that intact prophages are found in 88% of 192 published RSSC genomes, and that individual prophages are restrained to 2-3 continents and are phylotype-specific (50). Few exceptions are highlighted, one being phage Dina from Mauritius, a temperate phage with a high PHRI (generalist), which is distributed and abundant in multiple phylotypes from all six continents (50). As with CRISPRs, prophage prevalence is consistent with the PHRI data, and proves that generalism in Mauritian phages is a successful evolutionary strategy that assures widespread infections.

### 3. Bacteria-phage infection matrices are both modular and nested

To complete the host range analysis, we were interested in the structure of the phage-bacteria infection matrices and the possible identification of modules, composed of bacterial strains and phage species interacting specifically. In light of the different normalisations of titers and panels of bacteria tested, separate network analyses were performed for Mauritian and Reunion phages. A strong modularity pattern was detected in the Mauritius matrix, with relatively high modularity values (>0.35 on a scale spanning usually from 0 to 1, for all algorithms except *spinglass*) and a high statistical evidence (p-value <0.01) with all algorithms and all null models (Table 1, Fig. 1C). A weaker nested pattern was also detected, especially with the most efficient *WINE* algorithm and the most efficient C1+R1 and C2+R2 null models (Table 1), meaning that there is also a certain gradient of specialisation and generalism among phages. The analysis determined four modules in the Mauritian phage-bacteria matrix (Fig. 1C).

**Table 1:**
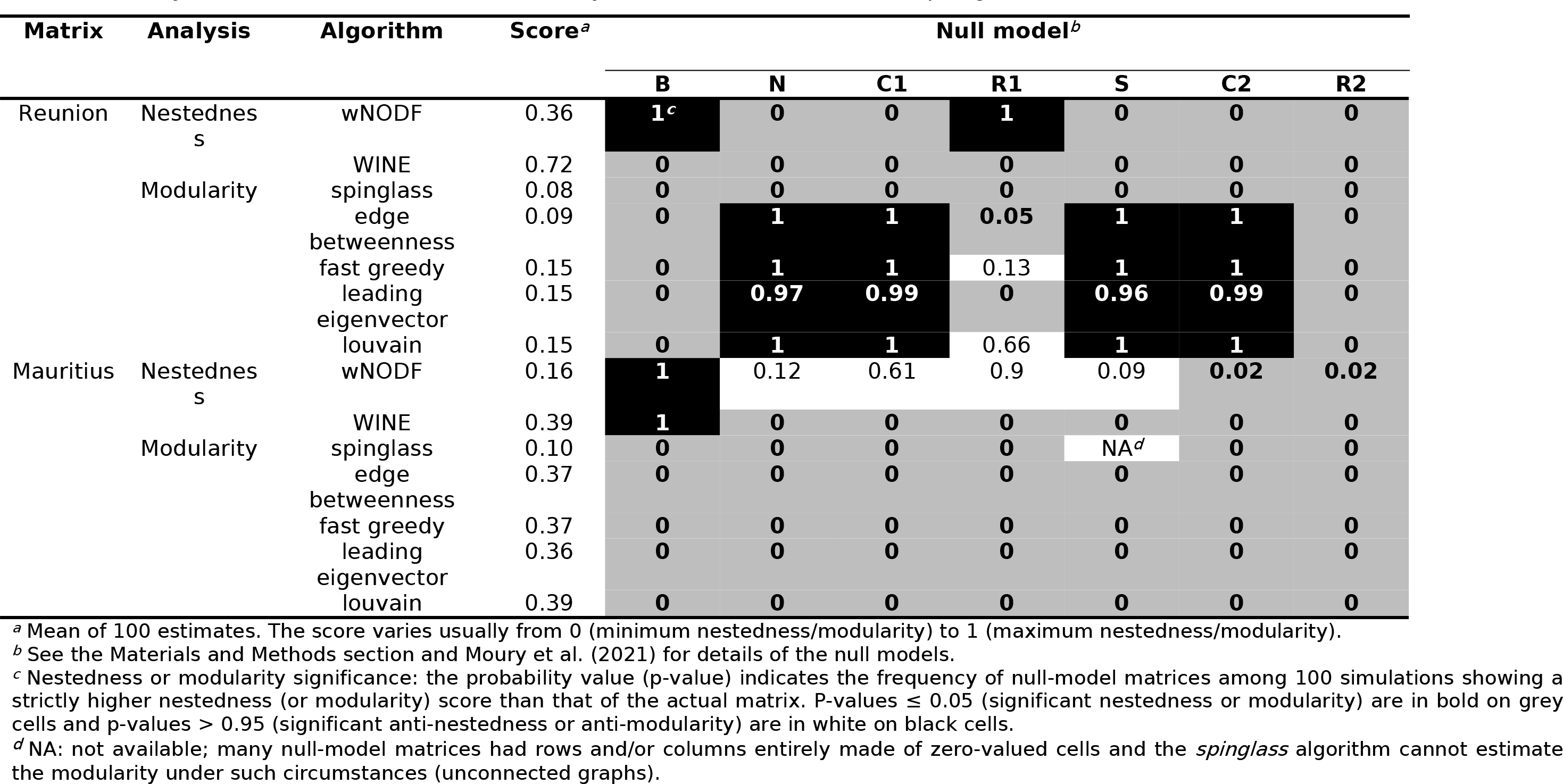
Analysis of nestnedness and modularity of Reunion and Mauritius phage-bacteria interaction matrices.

Module 1 was characterized by *Gervaisevirus Claudette* only, which is able to multiply in strains from phylotypes I and IV but not II. This phage is the only one targeting a particular strain of phylotype IV (RUN0062), isolated in Indonesia. Module 2 included six phages from different genera, such as *Firingavirus*, *Gervaisevirus* and *Dinavirus*, with the common feature that they can all target the same strain of sequevar I-15 (RUN4407), isolated in Mauritius. Module 3 integrated two Mauritian phages with a wide host range according to the PHRI analysis. Both phages targeted all three phylotypes but seemed specialised in strains from sequevar I-31 within phylotype I, similar to Reunion phages. These two phages belong to two different genera widely represented in Reunion, suggesting that they originated there and dispersed later to Mauritius. Phages of module 4 belong to the genera *Bakolyvirus* and *Cimandefvirus* and were able to target many strains from sequevar I-33, a few from other phylotype I strains and occasionally strains from phylotypes II and IV. These phages belong to genera also found in Reunion but, especially those from the genus *Bakolyvirus*, were extensively sampled in Mauritius Island (35). This implies that they could have first appeared in Mauritius and expanded later to Reunion Island, the opposite scenario described for the phages of module 3. At hindsight, modules are not always associated with phage taxonomy, with phages from the same genera and even same species belonging to different modules (Fig. 1C). This suggests a fast evolution of some phage genes, probably tail fibres recognising bacterial receptors, but not at a complete genomic level. This is not surprising, taking into account the high potential of mutation and recombination of phages (i.e., the genetic mosaicism of phages) (51, 52).

The phage-bacteria infection matrix from Reunion had a different structure. With the most efficient *WINE* algorithm, a high nestedness score was estimated (0.72) and all null models revealed a significant pattern (p-value < 0.01; Table 1), suggesting that, in spite of the homogeneous PHRI, there were both relatively specialist and generalist phages (Fig. 1D). Phages Anchaing, Cimandef and Dimitile were the relatively generalist ones, and Heva the most specialized, as confirmed by the previous PHRI analysis. A weak (0.08) but significant (p-value < 0.01) modularity score was also detected with the *spinglass* algorithm for all null models (Table 1). The four other modularity algorithms did not detect any significant modularity with the most efficient C1+R1 or C2+R2 null models. Similar to Mauritian phages, phage taxonomy was not always associated to the host range pattern, suggesting that local adaptation of phages is the most important explanatory variable. The nested structure revealed the most sensitive bacteria, namely sequevar I-31, and the more resistant ones, sequevars I-13 and I-15.

### 4. Phylogenetic signal of bacterial hosts and bacterial sensitivity differ between phages from the two islands

To test if the phage host range patterns detected were associated with a phylogenetic signal among the targeted bacteria, we calculated the genetic divergence of the strains infected by each phage. The phylogenetic signal of Mauritian phages is significantly lower than that of Reunion phages (χ2 = 11.635, df = 1, p ˂ 0.001), meaning that the strains they target are much less phylogenetically related (Fig. 2B). This implies that in the modules detected in the matrix analysis, bacteria are not taxonomically related and that similar molecular interactions may be at place in genetically distinct bacteria, probably implying analogous phage receptors or defence systems. The phylogenetic signal test supports the results obtained with the higher PHRI (generalism) obtained for Mauritian phages. The high phylogenetic signal of the bacteria targeted by Reunion phages corroborates that bacterial epidemiology is likely the most determinant factor of phage host range. Differences in the genetic diversity of the tested phages are not involved in the host range width, as it is high for both Reunion (four genera and five species) and Mauritius (six genera and eight species) [35].

Relationships between the bacterial resistance spectrum and the efficiency of their resistance to phages can reveal important evolutionary trade-offs. We therefore analysed, among bacterial strains, the correlations between the resistance spectrum, i.e. the number of phages that did not replicate in a given bacterial strain, and the average sensitivity to phages (excluding phages that did not replicate). For Mauritius Island’s bacteria panel, no significant relationship between resistance spectrum and sensitivity was observed (Pearson’s r = -0.155; p-value = 0.37; Fig. S1A). In contrast, for Reunion bacteria panel, the Pearson’s coefficient of correlation was significantly negative (r = - 0.695; p-value = 6.7e-04; Fig. S1B) and hence the stronger the resistance of bacteria, the broader their resistance spectrum, contrary to the trade-off expectations. This indicates that bacterial resistance mimics the phage host range observed and thus supports the close co-evolution of phage- bacteria communities in each island. In Reunion, phage specialisation in the dominant bacteria may be a cause or a consequence of the broad and strong phage resistance of secondary bacterial variants. For the multiple RSSC variants present in Mauritius, numerous specific adaptation pathways between phages and bacteria have likely occurred, producing a modular interaction structure.

### 5. Phages inhibit local bacteria efficiently

The bacterial inhibition efficacy over the three most abundant RSSC strains was lower on average for the phages from Mauritius (23% of inhibitory effect) that for those from Reunion (71% of inhibitory effect) (F1,187 = 154.7, p ˂ 0.001) (Fig. 3). However, this could be explained by the fact that the three strains tested for Reunion phages were genetically more similar (three haplotypes from sequevar I-31) than those tested for the Mauritian phages (three different sequevars). In fact, the virulence of Mauritian phages was more varied, with phages specialised at inhibiting efficiently only one of the three bacteria tested. This is confirmed by the lack of significant differences in overall virulence among phages but a significant phage-bacterial strain interaction (F_20,66_ = 46.09, p ˂ 0.001). Five out of the eleven Mauritian phages tested inhibited over 75% of bacterial growth of at least one strain, but six phages reached lower virulence values, suggesting that they could be specialised on other (not tested) strains (Fig. 3A). Most Reunion phages inhibited efficiently the three tested bacterial strains, with Anchaing and Raharianne phages having the lowest virulence (<50%) (Fig. 3B). Individual Reunion phages showed significant differences in their virulence on the three bacterial strains (F_9,80_ = 6.945, p ˂ 0.001). This result was unexpected on account of the genetical similarity of Reunion bacteria and the specialised host range of Reunion phages. In all, 12 out of the 21 phages had a high virulence capacity and nine phages had medium or low virulence effects on the bacteria tested.

**Fig. 3.**
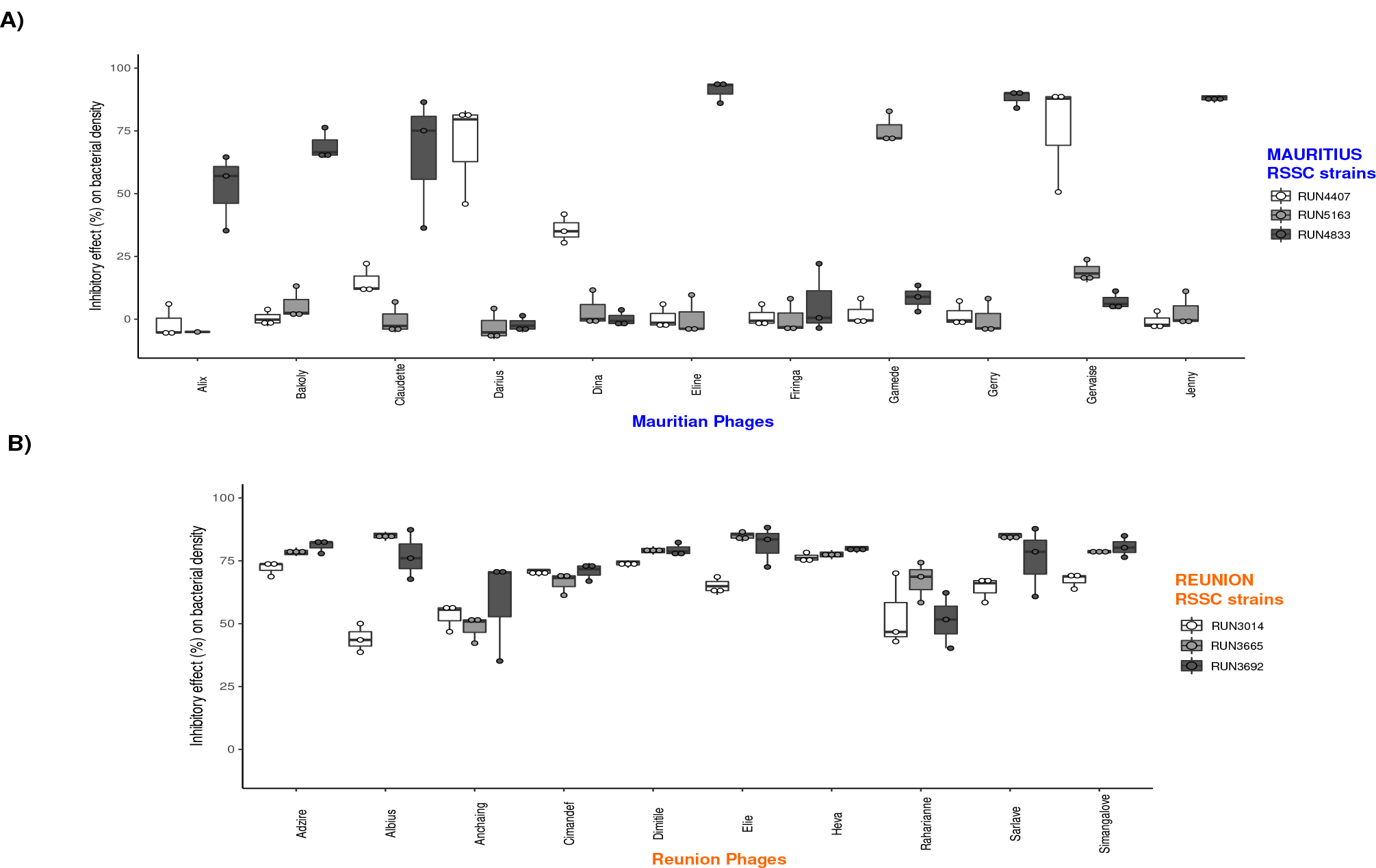
Virulence of phages measured as the inhibitory effect of phages on bacterial growth. Difference in mean bacterial density (OD600) over 24h in the presence and absence of phages. Eleven phages from Mauritius (A) and ten from Reunion (B) were analysed using the three most abundant RSSC strains on each island. Data are average phage virulence and standard errors.

The average fitness cost suffered upon phage attack in Mauritian RSSC variants differed significantly between sequevars I-33 (43%), I-15 (18%) and I-31 (9%) (F2,87 = 10.75, p ˂ 0.001) (Fig. 3A). However, no RSSC strain in Mauritius was broadly inhibited by the phages tested. Surprisingly, the strain RUN3014 did not suffer the highest fitness loss from infection by Reunion phages, in spite of the predominant abundance of this haplotype in Reunion. The variant RUN3665 was most inhibited (75%), followed by RUN3692 (73%) and lastly RUN3014 (64%) (F_2,87_ = 7.636, p ˂ 0.001) (Fig. 3B). The strong inhibition of strain RUN3665 is probably due to the fact that phages were propagated in this strain and may have subsequently adapted to it.

### 6. In complex epidemiology, generalist phages are more virulent; specialists thrive in simpler settings

We tested the hypothesis that generalist phages are less virulent than specialist ones, as a result of possible evolutionary trade-offs between host range breadth and virulence. No significant correlation between the PHRI and the maximum virulence value was observed for the whole dataset. A significant positive correlation was detected for Mauritian phages (Spearman’s ρ= 0.6818, df= 9, p = 0.0255) (Fig. 4). This result implies that the most virulent phages were able to target more RSSC strains, with a high genetic diversity. Also, it means that there is no apparent trade-off for generalist phages on their capacity to reduce the fitness of their hosts. For Reunion phages this trend was negative and significant, indicating that phage generalism comes at an evolutionary cost in a homogeneous bacterial epidemiology situation (Spearman’s ρ= -0.6727, df= 8, p = 0.0394) (Fig. 4).

**Fig. 4.**
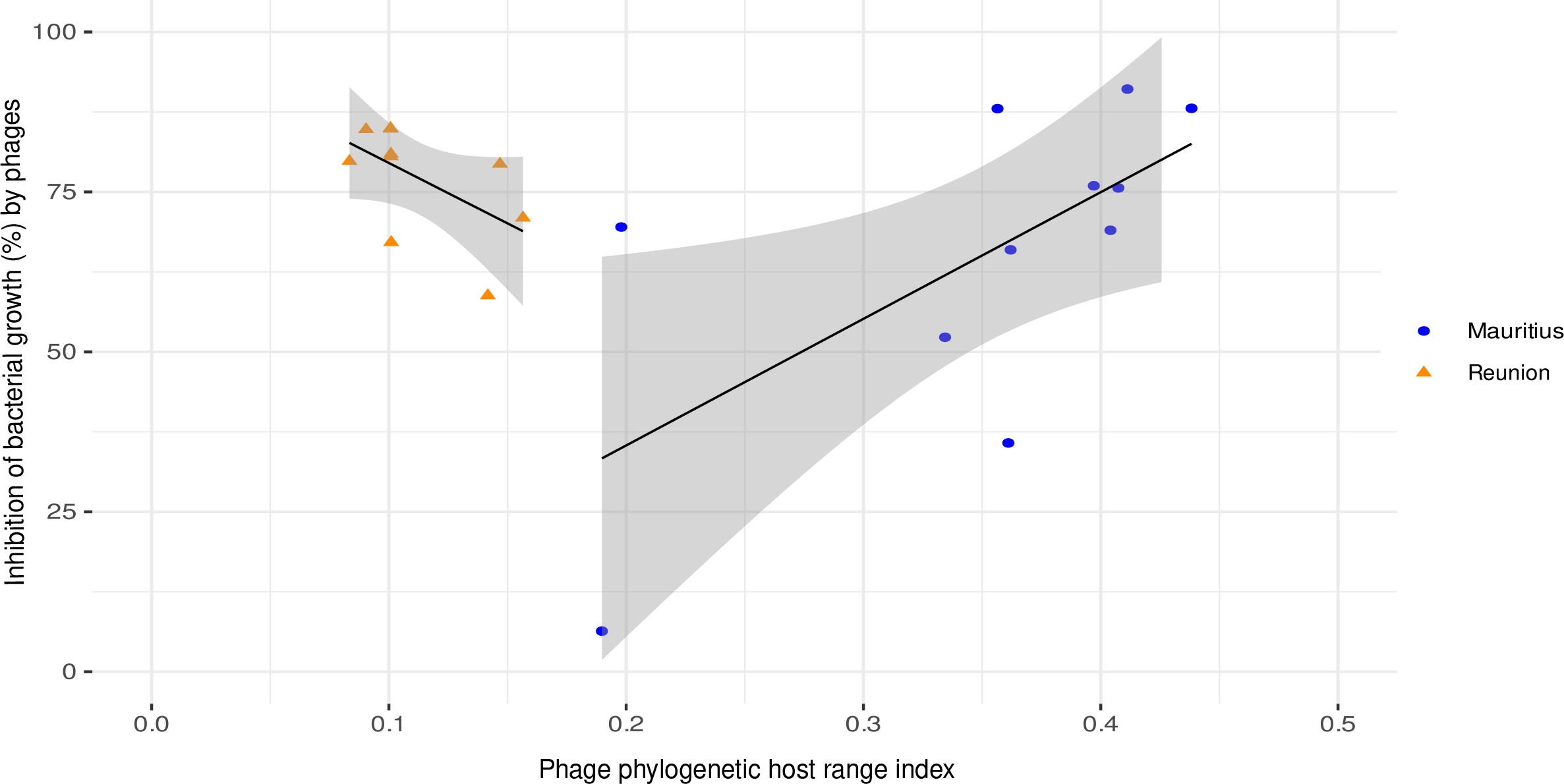
Correlation between maximum virulence, as the highest capacity of phages to inhibit bacterial growth, and the phylogenetic host range index of 21 phages, depending on the island of origin phages. Dataset of 13 Mauritian phages and 33 RSSC strains and ten Reunion phages, and 20 RSSC strains. Data are average values and the shaded area represents the standard error.

Positive correlations between these two traits have been proven in other organisms such as plant- feeding insects (53). We suggest that Mauritian phages may have optimised their replication rate and host inhibitory capacity as a consequence or as a prerequisite for their expanded host range. Reunion phages display a contrasted situation, where co-evolution between phages and bacteria in this island has selected for specialist phages. When the epidemiological landscape simplifies, specialists emerge as the most efficient phage type, presenting a precise adaptability to specific host environments. However, phages could display different virulence levels if other RSSC variants are tested, and we studied phages from only two locations attacking a particular bacterial species complex. Our study implies that the host range *vs* virulence evolutionary trade-off may not be common in phages and positive associations may be frequent in complex epidemiological situations. This conveys great advantages for generalist phage populations in nature and keys for the selection of candidate phages in therapeutic or biocontrol applied settings.

We also tested the association of phage virulence and host range with all phage variables available (genome size, protein content, GC%, taxonomy, etc). Most were non-significant (Table S3), but phage life cycle (virulent or temperate) was associated to phage virulence (χ2 = 6.6, df = 1, p = 0.0102) (Fig. S2), and to phage host range phylogenetic signal (χ2 = 6.7108, df = 1, p = 0.0096) (Fig. S3). Most phages with a maximum inhibitory capacity lower than 75 % are temperate. It has been described before that temperate phages usually have narrower host range and lower virulence (52). Even with the reduced number of temperate phages (6 out of 23) and high overall genetic diversity in our dataset, we detect this particular significant trend. The phage genome GC% was positively correlated to the PHRI (Spearman’s ρ= -0.4973, df= 23, p = 0.0158) (Fig. S4). The average GC% of all 23 phages is lower than that of the bacterial host, which has a median GC% of 66.8, pointing that phage niche expansion is related to adaptation to the host codon usage. Also, phage fitness was significantly associated with its morphology, with the highest for myovirus shapes (χ2 = 9.2977, df = 2, p = 0.0096) (Fig. S5), as previously shown (54).

Lastly, we investigated another crucial evolutionary trade-off within host-parasite interactions by testing whether the range of hosts a parasite can infect has an impact on its overall fitness. Among phages, the Pearson’s coefficient of correlation between PHRI and fitness was r = −0.802 for Reunion and r = 0.205 for Mauritius phages (Fig. S6). The r values obtained with random permutations of fitness values among phages ranged from −0.97 to 0.16 (mean −0.58) for Reunion and ranged from 0.021 to 0.395 (mean 0.208) for Mauritius ones. Out of the 1,000 random permutations, 61 and 486 yielded lower r values than those calculated for the actual fitness and host range data of Reunion and Mauritius phages, respectively. Therefore, the correlation between phage fitness and host range did not depart significantly from correlations obtained with random data permutations for Mauritius phages. For Reunion phages, the correlation was only slightly more negative than what was expected under the null hypothesis defined by random permutations, with a p-value of 0.061. (Fig. S6). Consequently, only a weak evidence of trade-off between the fitness and host range was obtained for phages from Reunion Island. Variables such as counter-defense mechanisms in phages and the characteristics of the receptor used, may play a role in determining the breadth of host range and its impact on fitness. We hope further research and more genomic data from both bacteria and phages, will shed light into this interesting topic.

## CONCLUSION

Host phylogenetic distance drives virus virulence and transmissibility (4). Phylogenetic information of the targeted species has been integrated for a long time in many ecological studies but only marginally to phage-bacteria ones (15, 17). Using an ecology phylogenetic diversity index to evaluate phage host range is a novel approach that provides with relevant information. We have determined the generalism or specialisation degrees of RSSC phages, which may reflect their past life history and epidemiological situation of the isolation place and may inform us about their future evolution. The enhanced matrix analysis applied to our phage-bacteria dataset complements this phylogenetic host range index. In ecology, modularity is regarded as an important feature for the maintenance of biodiversity, and the case of Mauritian phages and their modular interaction with RSSC bacterial variants supports that. Nonetheless, phage host range analyses are still limited by the frequent impossibility to integrate phage phylogeny when phages are very distant from each other (when no phylogenetic tree is possible or meaningful). Besides, an absolute phylogenetic phage host range index would be excellent to compare specialisation and generalism between phages tested against different bacterial strain panels. To achieve this, we sustain that laboratory and analytical methods of phage host range should be progressively standardized.

The association of phage host range with phage virulence has been poorly explored and we provide with evidence that there is an intricate interplay between phages and bacteria depending on the varying epidemiological complexity. Specialist phages are superior in more homogeneous ecological settings. Heterogeneous situations have likely led to a diversity of phage-bacteria interactions, including phage niche expansion, without paying apparent costs of being adapted to a wide range of hosts. However, much needs to be tested, as mutations that facilitate phage generalism and virulence may, in turn, lead to diminished fitness in different environments or evolutionary time scales (55, 56).

Our work relies on bacterial taxonomy, primarily focusing on a single gene. We questioned whether similar methods involving all available core-genes present in the core genome of bacteria, including phage receptor-coding genes, will produce with analogous results. CRISPR-Cas systems seem to play a role in defending bacteria against highly prevalent generalist phages, but the definite proof would be to generate genetically-engineered strains lacking these spacers and to undertake more bacterial comparative genomics. Moreover, this precise molecular system could partially explain some differences observed in phage virulence, as CRISPR blocks phage infection and replication less efficiently when the spacers are located far away in the spacer locus (57).

In practical terms, integrating these type of approaches will help optimizing potential biocontrol candidates and the durability of phage applications. *Ralstonia* spp. phages represent a promising biocontrol solution against the induced disease that is gradually being unravelled (33). Similarly to the PHRI steamed from classic community ecology, literature on insect trophic networks and their use in biocontrol have largely reflected on these questions and should be revisited by phage researchers (58). Our findings carry significance in elucidating the co-evolutionary processes that influence the ecological dynamics between phages and their hosts, offering insights that can guide the development of tailored phage-based approaches for targeting bacterial diseases.

## ACKNOWLEDGEMENTS

We thank the Plant Protection Platform (3P, IBISA) in Reunion Island. We are grateful to Enric Frago for fruitful discussions, Mirana Gauche and Emmanuelle Jousselin for advice on *Ralstonia* spp. phylogeny, and Ana Pineda for inspiring insight on mindful scientific writing. Authors declare no competing interests.

