## Supplementary material for "A phylogenetic host range index reveals contrasted relationships between phage virulence and specialisation"

List of content:

Supplementary Methods, Tables S1, S2 and S3, and Figures S1 to S6.

**Supplementary Methods**

Methods to estimate nestedness and modularity are described in Weitz et al. [1]. Two algorithms were available to estimate nestedness of matrices with quantitative data: the weighted nestedness metric based on overlap and decreasing filling (wNODF algorithm) [2] and the weighted-interaction nestedness estimator (WINE algorithm) [3]. In the R software, the ‘nested’ and ‘wine’ functions were used to estimate the wNODF and WINE scores, respectively. We used five different algorithms implemented into R software to estimate modularity: the spinglass [4–6], edge betweenness [4], fast greedy [7], leading eigenvector [8] and louvain [9] and  algorithms. Additional algorithms (label prop and walktrap) did not allow correct estimations of modularity (data not shown).

To determine the statistical significance of the nestedness or modularity of the phage-bacteria interaction matrices, the nestedness/modularity scores of the actual matrices derived from experimental data were compared to those of simulated null-model matrices that are not expected to possess any nested or modular pattern. Actual matrices were compared to matrices simulated under seven different null models [10]. Nestedness (or modularity) is significant if the actual matrix is more nested (or modular) than at least 95% of the matrices simulated under a given null model. Moury et al. [10] compared the performance (type I and type II error rates) of the nestedness and modularity algorithms and associated null models (Supplementary Methods 2 in Moury et al. 2021). For nestedness, a higher statistical power was observed for WINE than wNODF algorithm, whatever the null model. Consequently, we focused mainly on the results of the WINE algorithm. For modularity, the spinglass algorithm was by far the most efficient in terms of type I error rate, whatever the null model. For the four modularity algorithms, the type I error rate varied greatly depending on the null model. Also, null models C1, R1, C2 and R2 were the most suitable for both nestedness and modularity in terms of performance. Indeed, matrices showing significant patterns of nestedness or modularity with both models C1 and R1 (or both models C2 and R2) had the lowest type I error rates [10].

**Table S1**: List of Ralstonia solanacearum strains utilized for assessing the host range of phages from Mauritius and Reunion islands. Includes phylogenetic assignment, year and location of isolation, along with the corresponding *egl* sequence utilized for sequevar classification.

**Table S2**: Genomic content comparison of six phage pairs, illustrating the quantity and type of shared and unique proteins (highlighted in orange), alongside coding gene lengths and positions.

**Table S3**: Summary statistics for pair phage variables tested. Non-parametric Kruskall Wallis tests are used for factor variables, and Spearman correlation tests for numeric variables. P-values in bold indicate statistical significant tests and have been illustrated in figures. *NA* indicates impossible tests due to spurious correlations that have been analysed accordingly.

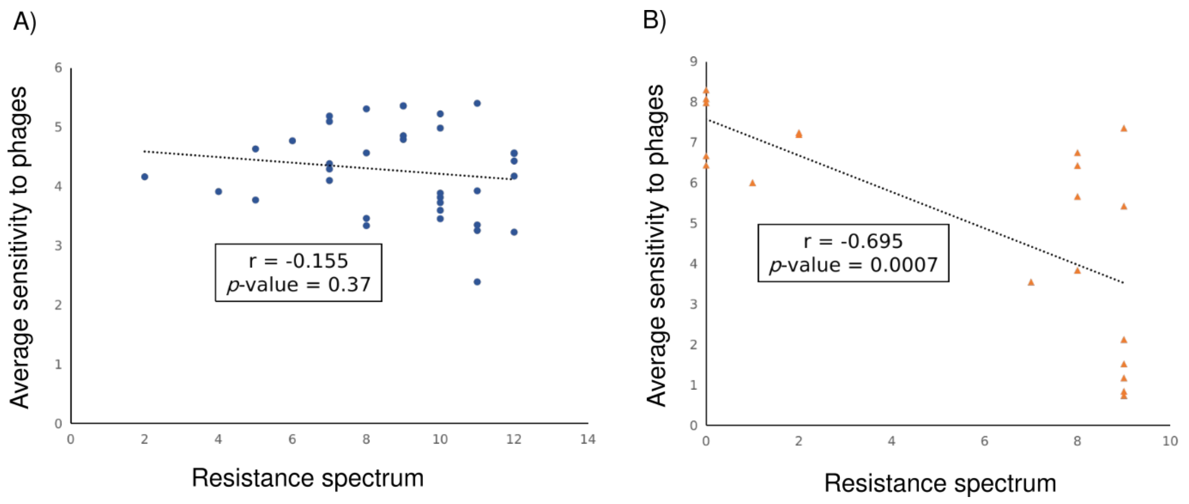


**Fig. S1**: Correlations between bacteria resistance spectrum and average sensitivity to phages. Thirty-six bacteria and 13 phages for Mauritius (A) and 20 bacteria and ten phages for Reunion (B) datasets.

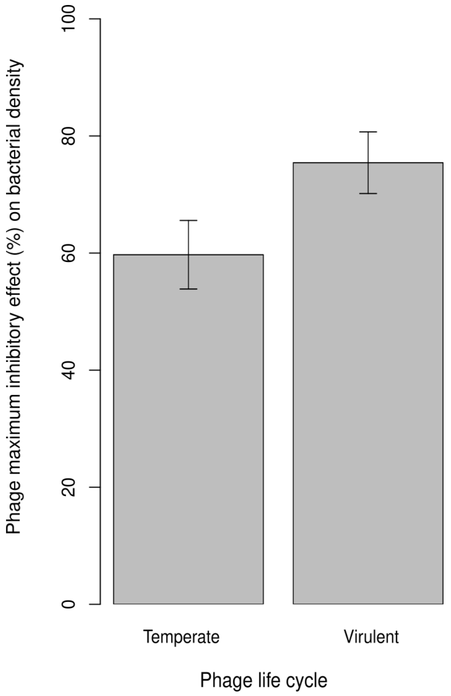


**Fig. S2:** Phage maximum inhibitory effect on bacterial density regarding the phage life cycle (temperate or virulent) of 21 phages.

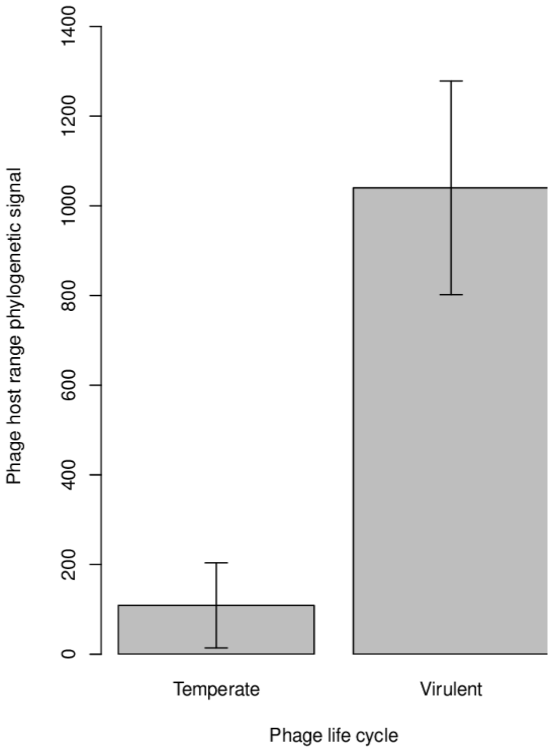


**Fig. S3**: Phage host range phylogenetic signal regarding the phage life cycle (temperate or virulent) of 23 phages.

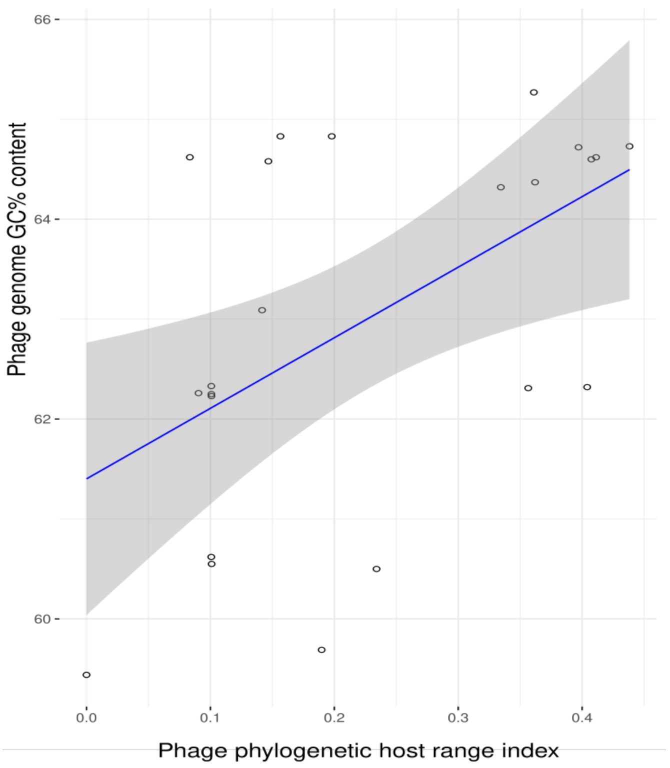


**Fig. S4:** Phylogenetic host range index relationship with the GC% genome content inhibitory effect in 23 phages.

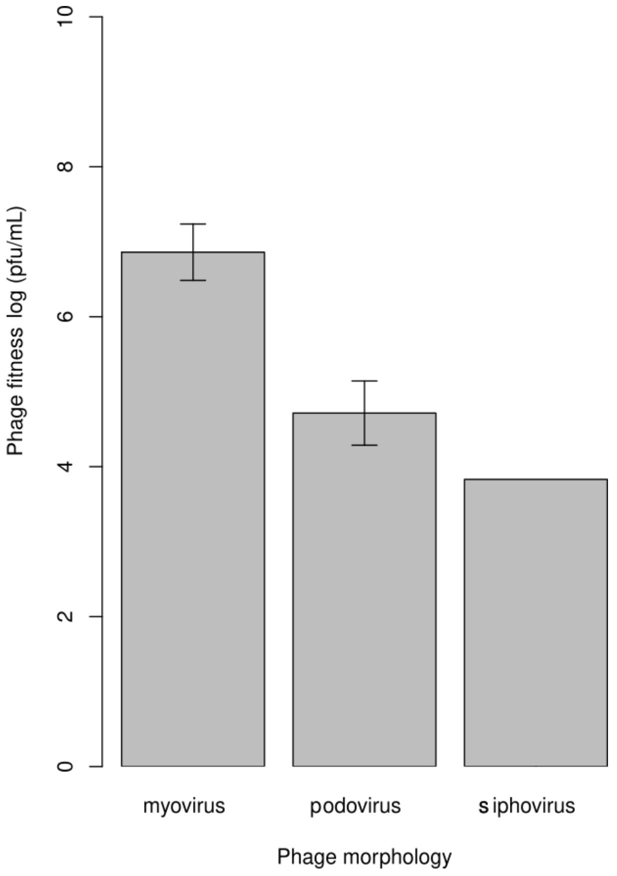


**Fig. S5**: Phage fitness regarding the phage morphology of 23 phages.

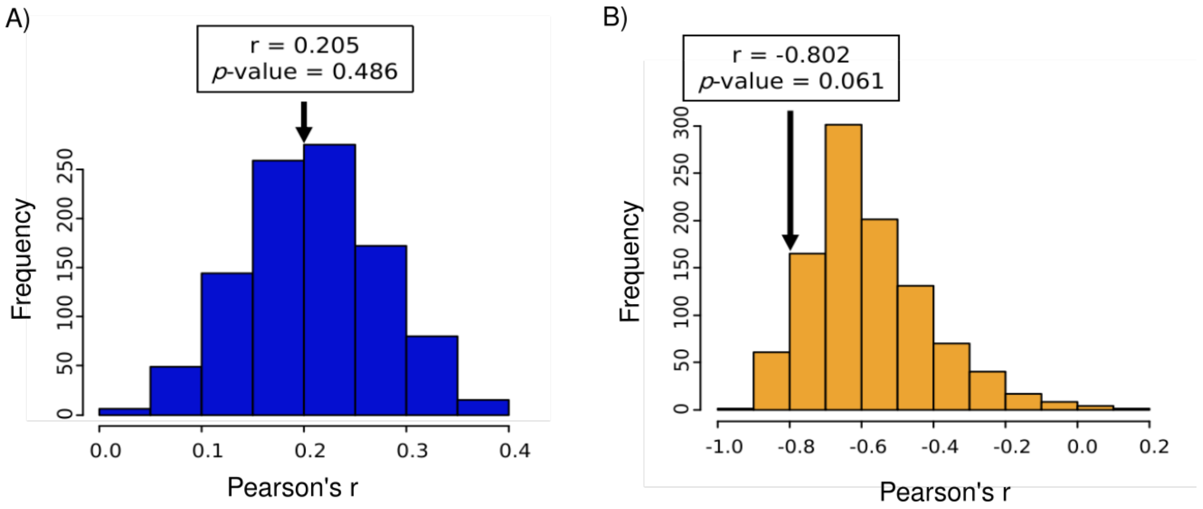


**Fig. S6:** Distribution of Pearson's r values between host range and fitness obtained using 1000 random permutations for phages from Mauritius (A) and Reunion (B) datasets. The actual r values are indicated by an arrow.
